# Cortical-brainstem interplay during speech perception in older adults with and without hearing loss

**DOI:** 10.1101/2022.06.03.494743

**Authors:** Jesyin Lai, Claude Alain, Gavin M. Bidelman

## Abstract

Real-time modulation of brainstem frequency-following responses (FFRs) by online changes in cortical arousal state via the corticofugal (top-down) pathway has been demonstrated previously in young adults and is more prominent in the presence of background noise. FFRs during high cortical arousal states also have a stronger relationship with speech perception. Aging is associated with increased auditory brain responses, which might reflect degraded inhibitory processing within the peripheral and ascending pathways, or changes in attentional control regulation via descending auditory pathways. Here, we tested the hypothesis that online corticofugal interplay is impacted by age-related hearing loss. We measured EEG in older adults with normal-hearing (NH) and mild to moderate hearing-loss (HL) while they performed speech identification tasks in different noise backgrounds. We measured α power to index online cortical arousal states during task engagement. Subsequently, we split brainstem speech-FFRs, on a trial-by-trial basis, according to fluctuations in concomitant cortical α power into low or high α FFRs to index cortical-brainstem modulation. We found cortical α power was smaller in the HL than NH group. In NH listeners, α-FFRs modulation for clear speech (i.e., without noise) also resembled that previously observed in younger adults for speech in noise. Cortical-brainstem modulation was further diminished in HL older adults in the clear condition and by noise in NH older adults. Machine learning classification showed low α FFR frequency spectra yielded higher accuracy for classifying listeners’ perceptual performance in both NH and HL participants. Moreover, low α FFRs decreased with increased hearing thresholds at 0.5-2 kHz for clear speech but noise generally reduced low α FFRs in the HL group. Collectively, our study reveals cortical arousal state actively shapes brainstem speech representations and provides a potential new mechanism for older listeners’ difficulties perceiving speech in cocktail party-like listening situations in the form of a miss-coordination between cortical and subcortical levels of auditory processing.

## 1 Introduction

Declines in auditory processing (Poth et al., 2001; Parthasarathy et al., 2010; Kortlang et al., 2016; Lai and Bartlett, 2018) and speech comprehension (Anderson et al., 2012) (Schneider et al., 2005; Peelle et al., 2010)—especially in the presence of background noise (Dubno, 1984; Takahashi and Bacon, 1992; Souza et al., 2007; Anderson et al., 2011; Song et al., 2011; Jin et al., 2014; Presacco et al., 2016; Vermeire et al., 2016)—are ubiquitous during aging and age-related hearing loss. Age-related declines in the sensory (auditory) system (Parthasarathy and Bartlett, 2011, 2012; Fostick et al., 2013; Parthasarathy et al., 2014, 2016; Lai and Bartlett, 2015, 2018; Lai et al., 2017)[e.g., age-related impairments in sound source segregation (Alain et al., 1996; Alain and McDonald, 2007; Gallun and Best, 2020)], changes in cognitive function (Park et al., 2003), or a combination of both (Pichora-Fuller and Singh, 2006; Wayne and Johnsrude, 2015; Wingfield et al., 2016) could lead to listening and comprehension difficulties in elderly listeners.

Evidence suggests that speech-in-noise (SiN) problems could be related to dysfunctional connections and changes in speech processing between cortical and subcortical levels of the auditory system that emerge with age and age-related hearing impairment. It is well established that SiN processing can be affected by many factors, such as attention (Saiz-Alía et al., 2019; Price and Bidelman, 2021) and arousal state (Mai et al., 2019; Saderi et al., 2021). Several studies have shown neural correlates of these phenomena. For example, findings from EEG studies on emotion suggest that power in the cortical α band (8-12 Hz) is a useful indicator of arousal state (Aftanas et al., 2002; Uusberg et al., 2013). Moreover, parieto-occipital α power was shown to index cognitive processing, effortful listening (Wöstmann et al., 2015; McMahon et al., 2016; Dimitrijevic et al., 2017), the state of wakefulness (Pfurtscheller et al., 1996) and top-down processing (Henry et al., 2017). Alpha oscillatory activity has also been associated with adaptive, intentional, and top-down suppression of task-irrelevant information (Rihs et al., 2007; Jensen and Mazaheri, 2010; Händel et al., 2011; Klatt et al., 2020). Increased α power has been proposed to index inhibitory processing across sensory modalities (Klimesch et al., 2007; Weisz et al., 2007, 2011; Strauß et al., 2014), while decreased α oscillations is thought to facilitate sensory processing or neural firing (Haegens et al., 2011; Klatt et al., 2020). There is, however, no consensus regarding the mechanisms underlying α oscillations reported in these studies. This could be due to the fact that in most studies α is treated as a unitary measure rather than reflecting different underlying processes. Meanwhile, evidence suggests that cortical α oscillations changes with aging (Yordanova et al., 1998; Böttger et al., 2002), such as a decrease in α frequency (Chiang et al., 2011) and reduced spontaneous entrainment of resting-state α oscillations (Gaál et al., 2010). Studying α power during SiN perception in older adults may reveal the impacts of aging in top-down attentional control that help facilitate the processing of target vs. non-target sounds, thus providing insight concerning why cocktail party-like situations are more difficult in older listeners (Pichora-Fuller et al., 2017).

In addition to cortical changes, age-related declines in speech coding have been widely observed at subcortical levels of the auditory system, both in terms of local processing within the brainstem but also its functional signaling to and from the cortex (Bidelman et al., 2019). In young adults, we recently observed that speech-evoked brainstem frequency-following response (FFR) amplitude varied as a function of α power (Lai et al, 2022). That is, low FFR amplitude coincided with low α power whereas high FFR amplitude was associated with high α states. Notably, low α FFRs correlated positively with behavioral response times (RTs) for speech discrimination and more accurately decoded the input speech stimuli as revealed by neural classifiers. Extending this approach here to address questions of auditory aging, we analyzed neuroelectric FFRs recorded during active speech perception in age-matched older adults with normal (NH) or mild hearing loss (HL). This allowed us to investigate the effects of age-related hearing loss on cortical α state and its modulation of brainstem speech processing in real-time. We aimed to determine the nature of auditory cortical-brainstem interplay in older adults, and more critically, whether such online corticofugal engagement during SiN listening is altered due to hearing loss, as suggested in prior work (Bidelman et al., 2019). Our collective results reveal that brainstem speech-FFRs are dynamically modulated by fluctuations in cortical α state in normal-hearing listeners but this cortical-subcortical interplay declines in age-related hearing loss.

## 2 Methods and Materials

### 2.1 Participants

Detailed information on participants, informed consent, and demographics are reported in our original report detailing age-related changes in the brainstem and cortical evoked potentials (Bidelman et al., 2019). New analyses reported herein examine online changes in FFRs as a function of the simultaneous cortical state. All participants had no reported history of neurological or psychiatric illness. Participants were aged between 52 to 75 (69 ± 5.8 years; 16/16 M/F). There were divided into normal (NH) and hearing-impaired (HL) groups based on their pure-tone audiometry hearing thresholds. We used 25 dB HL as the cutoff to define normal hearing, which has been used in the clinic since 1970s (Gatlin and Dhar, 2021). NH listeners (n = 13) listeners had average thresholds (250-8000 Hz) better than 25 dB HL across both ears whereas HL listeners (n = 19) had average thresholds poorer than 25 dB HL. The pure-tone averages (PTAs) (i.e., mean of 500, 1000, 2000 Hz) of NH listeners were ~10 dB better than in HL listeners (mean ± SD; NH: 15.3 ± 3.27 dB HL, HL: 26.4 ± 7.1 dB HL; t_2.71_ = −5.95, p < 0.0001; NH range = 8.3-20.83 dB HL, HL range = 15.8-45 dB HL). Both NH (t_12_ = 0.15, p = 0.89) and HL (t_18_ = −2.02, p = 0.06) groups otherwise had symmetric PTA between ears. Both NH and HL groups had elevated hearing thresholds at very high frequencies (≥ 8000 Hz), which is typical of age-related presbycusis in older adults. Besides hearing, the two groups were matched in age (NH: 66.2 ± 6.1 years, HL: 70.4 ± 4.9 years; t_2.22_ =−2.05, p = 0.052) and sex balance (NH: 5/8 M/F; HL: 11/8; Fisher's exact test, p = 0.47). Age and hearing loss were not correlated (Pearson's r = 0.29, p = 0.10), suggesting these factors of aging were largely independent in our sample.

### 2.2 QuickSiN Test

The Quick Speech-in-Noise (QuickSiN) test was used to measure listeners’ speech reception thresholds in noise (Killion et al., 2004). A list of six sentences with five keywords per sentence spoken by a female talker in a background of four-talker babble noise was heard by listeners during the test. Target sentences were presented at 70 dB sound pressure level (SPL) (binaurally) at signal-to-noise ratios (SNRs) decreasing from 25 dB (relatively easy) to 0 dB (relatively difficult) in 5 dB steps. The number of keywords correctly recalled was logged, and a score was computed for each listener. The SNR-loss score indexes the difference between a listener’s SNR-50 (i.e., the SNR required to identify 50 % of the keywords correctly) and the average SNR threshold for normal-hearing adults (i.e., 2 dB) (Killion et al., 2004). A higher score reflects poorer SiN perception. Each listener’s SNR-loss score was averaged from four lists of sentence presentations. In this study, NH listeners' scores ranged from −0.25 to 2.5 dB of SNR-loss (M = 1.1, SD = 0.8) while HL listeners’ scores ranged from −2.5 to 8.5 of SNR-loss (M = 2, SD= 2.5) (see Fig. 1D in Bidelman et al., 2019).

**Figure 1.**
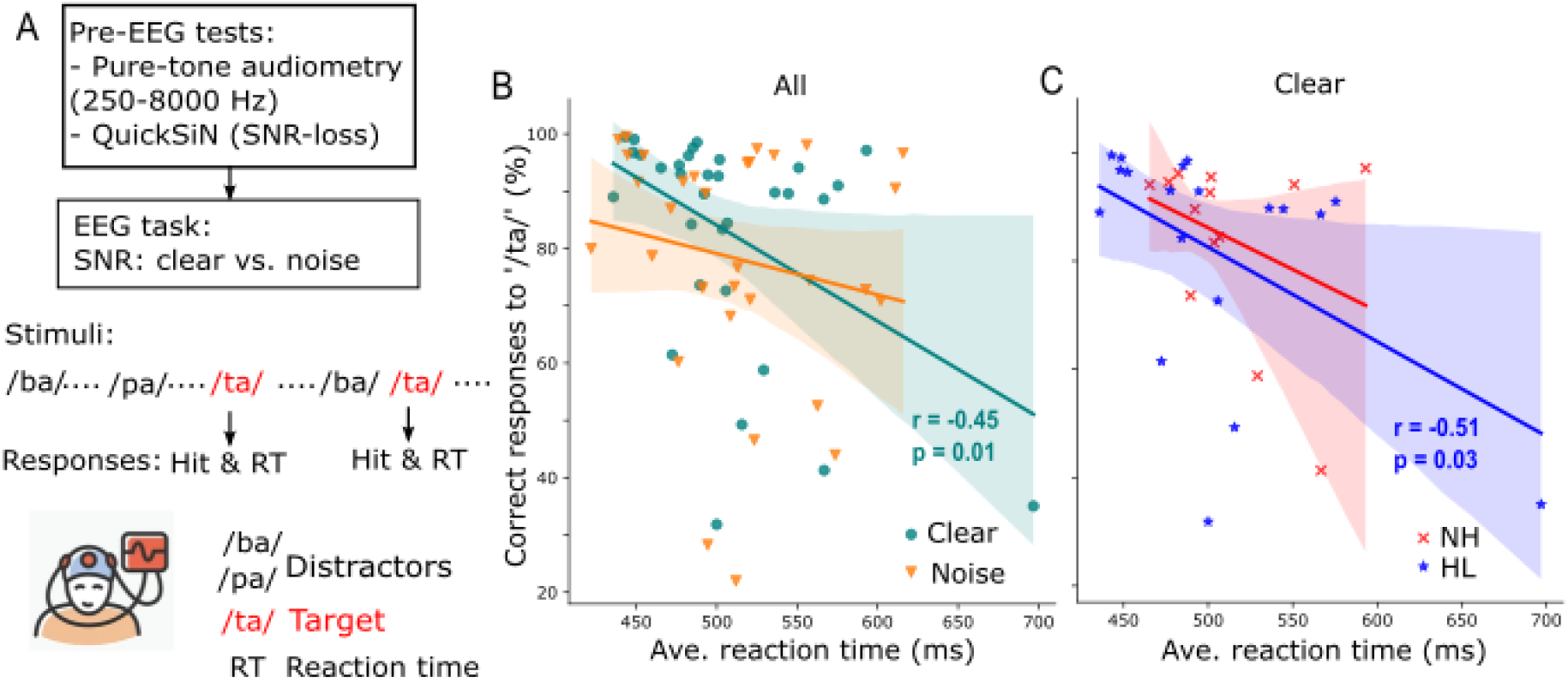
Target speech detection performance correlates with average response time collected during an active EEG task. (A) Prior to EEG recordings, all participants' pure-tone audiometry tested at 250-8000 Hz were obtained and speech-in-noise perception was assessed with QuickSiN. Subsequently, speech-EEGs to consonant-vowel phonemes (/ba/, /pa/, and /ta/) were recorded under clear or noisy (+10 dB SNR) backgrounds while participants actively engaged in the target speech detection task. (B) Correct responses to target speech (i.e., /ta/) are predictive of reaction times in quite (clear) but not noisy backgrounds. (C) When participants were divided into normal (NH) and hearing loss (HL) groups, the correlation of speech detection accuracy and reaction time obtained in quiet backgrounds is significant only in the HL group. r = Spearman's correlation; shaded area = 95% CI of the regression line.

### 2.3 EEG Stimuli and Task

The stimuli and task are described fully in Bidelman et al. (2019) and illustrated here in Fig. 1A. Three naturally produced English consonant-vowel phonemes (/ba/, /pa/ and /ta/), from the standardized UCLA version of the Nonsense Syllable Test (Dubno and Schaefer, 1992), were generated by a female talker. The duration of each phoneme was 100-ms and the average root mean square SPL of each phoneme matched. All three tokens had a common voice fundamental frequency (mean F0 = 150 Hz), first and second formants (F1 = 885, F2 = 1389 Hz). The resulting stimulus-evoked response (i.e., FFR) should predominantly originate from the subcortex (Brugge et al., 2009; Bidelman, 2018) since the stimulus F0 is above the phase-locking limit of the cortical neurons and “cortical FFRs” (Coffey et al., 2016; Bidelman, 2018; Bidelman and Momtaz, 2021). Speech tokens were delivered binaurally to listeners in either clear (i.e., no noise) or noise-degraded conditions. A complete set of stimulus presentation in each condition contained a total of 3000 /ba/, 3000 /pa/, and 210 /ta/tokens (spread evenly over 3 blocks to allow for breaks). The interstimulus interval between tokens was randomly jittered within the range of 95-155 ms (5ms steps, uniform distribution). The /ba/ and /pa/ tokens were presented more frequently than the /ta/ token in a pseudo-random manner such that at least two frequent tokens intervened between infrequent tokens. The infrequent /ta/ token was denoted as the target in which listeners were required to provide a response by pressing a button on the computer whenever they detected it. Both reaction time (RT) and detection accuracy (%) were recorded. For the noise-degraded condition, the same procedures as the clear condition were repeated, but the tokens were presented in an identical speech triplet mixed with eight talker noise babble (Killion et al., 2004) at a signal-to-noise ratio (SNR) of 10 dB. There were 6 blocks (3 clear & 3 noise) collected from each participant. Having the clear and noise conditions allowed us to compare behavioral performance in different backgrounds and evaluate the impact of noise on speech perception in NH vs. HL listeners, respectively. The task ensured that listeners were actively engaged during speech perception and online EEG recording. Stimuli were controlled by a MATLAB program (The Mathworks, Inc.; Natick, MA) routed to a TDT RP2 interface (Tucker-Davis Technologies; Alachua, FL) and delivered binaurally through insert earphones (ER-3; Etymotic Research; Elk Grove Village, IL). The speech stimuli were presented at 75 dB SPL (noise at 65 dB SPL) with alternating polarity.

### 2.4 EEG recording and preprocessing

The neuroelectric activity was recorded from 32 channels at standard 10-20 electrode locations on the scalp (Oostenveld and Praamstra, 2001) during the target speech detection task. Electrode impedances were ≤ 5 kΩ. EEGs were digitized at 20 kHz using SynAmps RT amplifiers (Compumedics Neuroscan; Charlotte, NC). After EEG acquisition, the data were processed using the ‘mne’ package in Python 3.9.7. EEG data were re-referenced offline to the mastoids (TP9/10) for sensor (channel-level) analyses. For source analysis of brainstem FFRs, we used a common average reference prior to source transformation (detailed below). Responses were then filtered 100-1000 Hz [finite impulse response (FIR) filters; hamming window with 0.02 dB passband ripple, 53 dB stopband attenuation, −6 dB cutoff frequency] to further isolate brainstem activity (Musacchia et al., 2008; Bidelman et al., 2013).

### 2.5 Derivation of Source FFRs and Cortical Activities

The derivation of source FFR waveforms and isolation of cortical activities are similar to the methods described in Lai et al. (2022) for young adults. The 32-ch sensor data were transformed into source space using a virtual source montage. The source montage comprised of a single regional source (i.e., current flow in x, y, z planes) positioned in the brainstem and midbrain (i.e., inferior colliculus) [details refer to (Bidelman, 2018b; Bidelman and Momtaz, 2021; Price and Bidelman, 2021)]. Source current waveforms (SWF) from the brainstem source were obtained using the formula: SWF = L^−1^ × FFR, where L is the brainstem source leadfield matrix (size 3×64) and FFR is the 32-ch sensor data (64 × NSamples). This essentially applied a spatial filter to all electrodes that calculated their weighted contribution to the scalp-recorded FFRs in order to estimate source activity within the midbrain in the x, y, and z directions (Scherg and Ebersole, 1994; Scherg et al., 2002). This model explains ~90 % of the scalp-recorded FFR (Bidelman et al., 2019; Price and Bidelman, 2021). Only the z-oriented SWF was used for further analysis (x and y SWFs were not analyzed) given the predominantly vertical orientation of current flow in the auditory midbrain pathways relative to the scalp [x- and y-orientations contribute little to the FFR (Bidelman, 2018b)].

We isolated cortical α band activity from the EEG and used it as a running index of arousal state (high or low) during the target speech detection task. EEG at the Pz and Oz channels were filtered at 8-12 Hz (FIR filters, −6 dB cutoff frequency at 7 Hz and 13.5 Hz) and averaged (i.e., equivalent to POz) to obtain cortical α-band activity at a posterior scalp region. Filtered α activities were epoched with a time window of 195 ms (−50 to 145 ms in which 0 ms corresponded to the onset of a /ba/ or /pa/ token) to capture approximately 1-2 cycles of α band. This epoch window encapsulated the entirety of the evoked FFR within the immediate trial with no spillover from the preceding or subsequent trial(s). Infrequent /ta/ tokens were excluded from analysis due to their limited trials. The root mean square (RMS) amplitude of single trial α activity was computed to quantify cortical arousal level over the duration of the target speech detection task. We then normalized RMS values to the median of RMS values of each run, respectively. Next, the distribution of trial-wise normalized α RMS was visualized using a histogram. We categorized trials of each participant per condition falling within the 0-35th percentile as “low α” power and those falling within the 65-100th percentile as “high α” power. This categorization was used because it provided ~2100 trials for each low or high α power in each participant per condition, which is reasonable to obtain an average FFR with robust signal-to -noise ratio (i.e., ≥3 dB SNR) (Bidelman, 2018a). More detailed information of this methodology can be found in Lai et al. (2022) (see their Fig. 2). We similarly measured cortical activity in another frequency band (e.g., β band; 18-22 Hz) from the same location (i.e., POz β) and α band from a different electrode site (i.e., Fz α). These control analyses allowed us to ensure that the observed changes in speech-evoked FFRs were specifically associated with cortical arousal level (indexed by *α* power) rather than general fluctuations in the EEG, per se.

**Figure 2.**
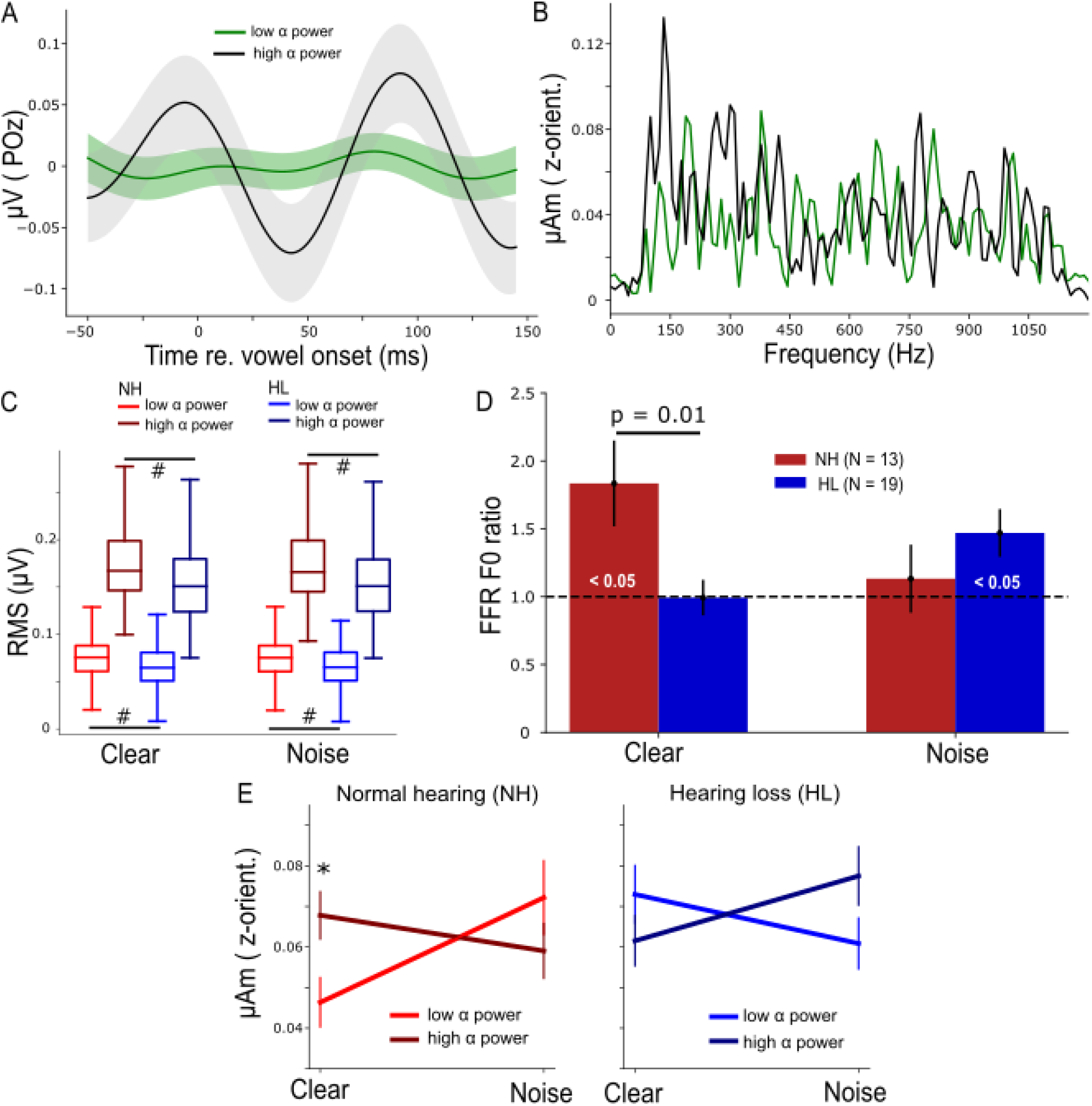
Normal hearing (NH) participants have a higher parieto-occipital α power and their brainstem speech processing is more strongly modulated by α band in clear backgrounds. (A) Average α waveform of low- and high-power trials plotted from a representative NH participant. (B) Frequency spectra of the steady state (10-100 ms) portion of low and high α brainstem FFRs from the same participant in (A). Note the distinct response at ~150 Hz, corresponding to the voice pitch. (C) Root-mean-square (RMS) values of both low and high α of the NH group were significantly higher than the HL group. # p< 0.01 (Conover’s test, non-parametric pairwise test, with Bonferroni adjustment) (D) FFR F0 ratios during low and high α trials. FFR F0 ratios were higher in the NH vs. HL group (Mann-Whitney U test) in the clear condition. Bars marked (<0.05) are significantly larger than 1 (1-sample t-test) indicating enhancement of the FFR with changes in cortical α. (E) Grand average FFR F0 amplitudes as a function of SNR (clear vs. noise), α power (low vs. high), and group (NH vs. HL). Error bars= ± s.e.m., *p<0.05 (Wilcoxon signed-rank test).

### 2.6 Analysis of Brainstem FFRs

We categorized source FFRs based on whether α amplitude in the same epoch was either high or low power, thus deriving FFRs according to the trial-by-trial cortical state. Source FFR waveforms (from the z-orientated dipole) were averaged for each α category and noise condition per participant. Subsequently, we analyzed the steady-state portion (10–100 ms) of FFR waveforms using the FFT (Blackman window; 11.1 Hz frequency resolution) to capture the spectral composition of the response. F0 amplitude was measured as the peak spectral maximum within an 11 Hz bin centered around 150 Hz (i.e., F0 of the stimuli). To compare FFR F0 amplitudes during low vs. high α power, a normalized (within-subject) F0 ratio was calculated as follows:

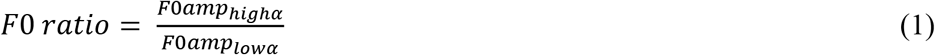

Where F0 ratios > 1 indicate stronger brainstem FFRs during states of high cortical α power and F0 ratios < 1 indicate stronger FFRs during states of low cortical α power (Lai et al., 2022).

### 2.7 Statistical Analysis

We used rmANOVAs to compare brainstem F0 ratios among the clear vs. noise condition, and NH vs. HL group. Multiple pairwise comparisons (Mann-Whitney U test with Bonferonni corrections) between the NH and HL groups were performed using the ‘pingouin’ package in Python. One sample t-tests (‘scipy’ package in Python) were also used to evaluate whether FFR F0 ratios were significantly different from 1 (and thereby indicated the significance of α modulation). Wilcoxon signed-rank test was used when comparing raw F0 amplitudes at low vs. high α power within participants for each SNR condition and hearing group. To compare differences in α RMS values of all participants across (clear vs. noise) conditions, we performed post-hoc Conover’s test (“scikit_posthocs” package in Python), which is a non-parametric pairwise test, with Bonferroni adjustment. To assess differences in raw F0 amplitudes (log-transformed) across factors for NH and HL groups, we first performed a 2 × 2 × 2 (SNR × α power × hearing group) mixed model (participants = random factor) ANOVA (‘lme4’ package in Rstudio). Following a significant interaction, we then ran separate 2 × 2 (α power × SNR) mixed-model ANOVAs for the NH and HL groups, respectively. Initial diagnostics were performed using residual and Q-Q plots to assess the heteroscedasticity and normality of data. F0 amplitudes were log-transformed to improve normality and homogeneity of variance assumptions. Effect sizes are reported as η_p_^2^. To check if low or high α FFR F0 amplitudes were associated with behavioral performance (i.e., QuickSiN, PTA, %-correct, and RTs) by pooling all NH and HL participants, we performed Spearman’s correlations (‘scipy’ package in Python) to test their pairwise correlations of brain and behavior measures. Spearman’s correlation was used because these measures were found to be not normally distributed (p < 0.05) from the test of normality using Shapiro-Wilk test.

### 2.8 Classification of Performance Level from FFR Frequency Spectra via Machine Learning

All participants’ performance in the clear condition was categorized into three levels (poor, average, and good) based on their percent correct of /ta/ detections. Participants with poor performance had %-correct ≤ 30th-percentile of overall %-correct while good-performance participants had %-correct ≥ 70th-percentile of overall %-correct. Subsequently, we classified poor- and good-performing participants using frequency spectra of their low or high α FFRs and a support vector machine (SVM) classifier (kernel = radius basis function, C=1000, gamma = scale) in the ‘scikit-learn’ package in python. Due to the limitation in sample size (32 total observations), we were only able to perform a two-group rather than three-group machine learning (ML) classification. There were ten participants falling into either poor- or good-performance category, which provided twenty participants’ frequency spectra to be used as data input for SVM. Frequency spectra were obtained from the FFT of average FFR waveforms (10 to 100 ms steady state portion) across ~2100 trials of low or high α FFRs per participant. The absolute amplitudes of frequency spectra, which consisted of 226 amplitude-by-frequency points, were the input features; performance level (i.e., poor vs. good) served as the ground truth class labels. The ML classification procedures on FFRs were similar to those described by Xie et al. (2009) and Lai et al. (2022). During one iteration of training and testing, a four-fold cross-validation approach was used to train and evaluate the performance of the SVM classifier to obtain a mean classification accuracy [Fig. 1 of Xie et al. (2009)]. In this process, a 4-fold stratification was performed to randomly and equally divide participants into 4 subgroups with 5 unique participants (almost similar number of poor- and good-performing participants) in each subgroup. Three of the 4 subgroups were selected as the training data while the remaining subgroup was used as the hold-out testing data. To mitigate the problem of imbalanced participant numbers in the two classes of training set, we randomly over-sampled the minority class using the ‘imblearn’ package in python. These steps were repeated within each iteration so that each subgroup was held-out as the test data whereas the other 3 subgroups were used to train the SVM classifier. Mean classification accuracy of poor- vs. good-performance was calculated across cross-validated iterations. We performed a total of N=5000 iterations to examine group classification for low vs. high α power FFRs.

To evaluate if the classifier accuracy (mean of N=5000 iterations) was statistically significant, we randomly shuffled the 226 data points of frequency spectra in each participant, and the same training and testing procedures described above were repeated to derive a null distribution of classification accuracies. We then calculated the p-value to determine the statistical significance of "true" classifier performance using the formula described in Phipson and Smyth (2010): *p = (a + 1)/(n + 1)*, where a is the number of classification accuracies from the null distribution that exceeds the median of the actual distribution of classification accuracies and n is the total number of classification accuracies from the null distribution.

### 2.9 Fitting Linear Regression Models with Brain and Behavior Measures

To compare the changes in low α FFR F0 amplitudes as behavior performance changed in the NH and HL groups, we fitted linear regression models for pooled SNR, clear and noise conditions, respectively, by using behavior performance and group as main factors, and their interaction factor.

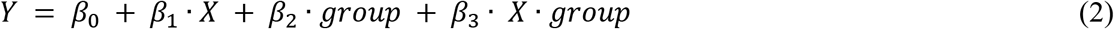

*Y* represents FFR F0 amplitudes during low α power, *X* is one of the behavior measures (QuickSiN, PTA, %-correct, or RTs), and *group* is the dummy variable for NH (*group=0*) and HL (*group=1*). Meanwhile, *β_0_* is a constant and *β_1_*, *β_2_*, or *β_3_* is the coefficient or slope for the respective variable. After fitting regression models, we checked assumptions for the normality of residuals, homoscedasticity of residuals and linearity of the models. For residual normality, we tested if model residuals were normally distributed using Shapiro-Wilk test and visualized the Q-Q plots of the residuals. For residual homoscedasticity, we used Breusch-Pagan and Goldfeld-Quandt tests. For model linearity, we visualized residual distributions by plotting residuals vs. predicted values. Result interpretation was then carried out after confirming the above three assumptions.

The fitted linear regression models allowed us to study if the slopes (i.e., the change of FFR amplitudes) of the NH group was significant as well as if the slope of the HL group was significantly different from the NH group. In the NH group, i.e., *group*=0, Eq. (2) can be written as *Y = β_0_ + β_1_*∙*X* and *β_1_* is the slope for the NH group. In contrast, in the HL group, i.e., *group*=1, Eq. (2) can be written as *Y = β_0_ + β_1_*∙*X+ β_2_+ β_3_*∙*X =* (*β_0_ + β_2_)* + (*β_1_+ β_3_)X*, where (*β_1_+ β_3_)* is the slope for the HL group and *β_3_* represents the difference in slope between the HL and NH group.

## 3 Results

### 3.1 Behavior performance of target speech detection

Behavioral responses during the EEG task (%-correct /ta/ detections vs. RTs) showed a negative correlation for the clear but not noise condition (Spearman’s r = −0.45, p = 0.01, Fig. 1B); participants with slower response speeds showed poorer speech detection accuracies. This is consistent with previous findings showing negative associations between hit responses and RTs in younger listeners (e.g., Lai et al. 2022). When separated into the NH and HL groups, we found a negative relationship between behavioral hit responses and decision speeds but only in the HL group (Fig. 1C).

### 3.2 Cortical α band and brainstem speech-FFRs

Differences in cortical α-band amplitudes during low vs. high α states were prominent at the single participant (Fig. 2A) as well as group level (Fig. 2C). Spectral differences in the corresponding brainstem FFR for these same low vs. high cortical trials were also notable (Fig. 2B). Cortical α (both low and high levels) was overall higher in the NH listeners (p<0.01, non-parametric post-hoc Conover’s test with Bonferroni adjustment) but both groups showed clear separability of “low” vs. “high” α states during the speech detection task.

In response to clear speech (Fig. 2D), brainstem F0 ratios (indexing cortical α-related FFR enhancement) in the NH group were significantly higher than 1 (t_12_ = 2.64, p = 0.02, 1-sample t-test) and higher than the HL group overall (U = 192, p = 0.01, Mann-Whitney U test). In response to the noise-degraded speech, this cortical-FFR enhancement was observed in the HL group (t_18_ = 2.67, p = 0.02, 1-sample t-test), but was not significantly different than the FFR enhancement observed in the NH group. Repeating the same analysis for both controls (POz β and Fz α) revealed no difference in F0 ratios for the NH vs. HL group (Fig. 1 in Supplementary), indicating cortical modulation of the FFR was restricted to POz α.

A 3-way mixed-model ANOVA performed on log F0 amplitudes revealed a significant SNR × α power × group interaction (F_1, 96_ = 9.5, p = 0.003, η_p_^2^ = 0.09) (Fig. 2E). To make sense of this complex interaction, we performed separate 2-way (SNR × α power) mixed-model ANOVAs by hearing group. The SNR × α power interaction was significant in both the NH (F_1,39_ = 5.17, p = 0.03, η_p_^2^ = 0.12) as well as the HL group (F_1,57_ = 4.11, p = 0.05, η_p_^2^ = 0.07). Though the comparison of effect sizes suggests this interaction was stronger in NH listeners, the interaction was distinct in direction compared to the HL group. In the NH group, FFR F0 amplitudes were significantly higher during high α power for clear speech. This pattern was dampened and reversed in the HL group.

### 3.3 Brain-behavior relations in both the NH and HL groups

We next assessed associations between low or high α FFR F0 amplitudes) and behavioral performance (QuickSiN, PTA, %-correct, and RTs) by performing Spearman’s correlation analyses. The positive association between F0 amplitudes and percent correct was significant at low but not high α power (Spearman’s r = 0.3, p = 0.02, Fig. 3A) while the associations between F0 amplitudes (either during low or high α power) and other behavior measures were not significant. To further assess if FFRs during low or high α power in either clear or noise condition were more predictive of behavioral responses, we performed ML classification of percent correct (or perceptual performance level) into poor (≤ 30th of overall percent correct) or good (≥ 70th of overall percent correct) using participants’ FFR frequency spectra as input for the SVM classifiers. In the clear condition (Fig. 3B), the classifier was significantly better in decoding participants’ perceptual performance level using low α FFRs compared to high α FFRs. In contrast, in the noise condition (Fig. 3C), classification accuracies did not differ between low or high α FFRs nor where they above the null distribution. These results provide evidence that adding background noise disrupted the relationship of low α FFRs with behavioral measures potentially as a result of compromising the SNR of the neural responses.

**Figure 3.**
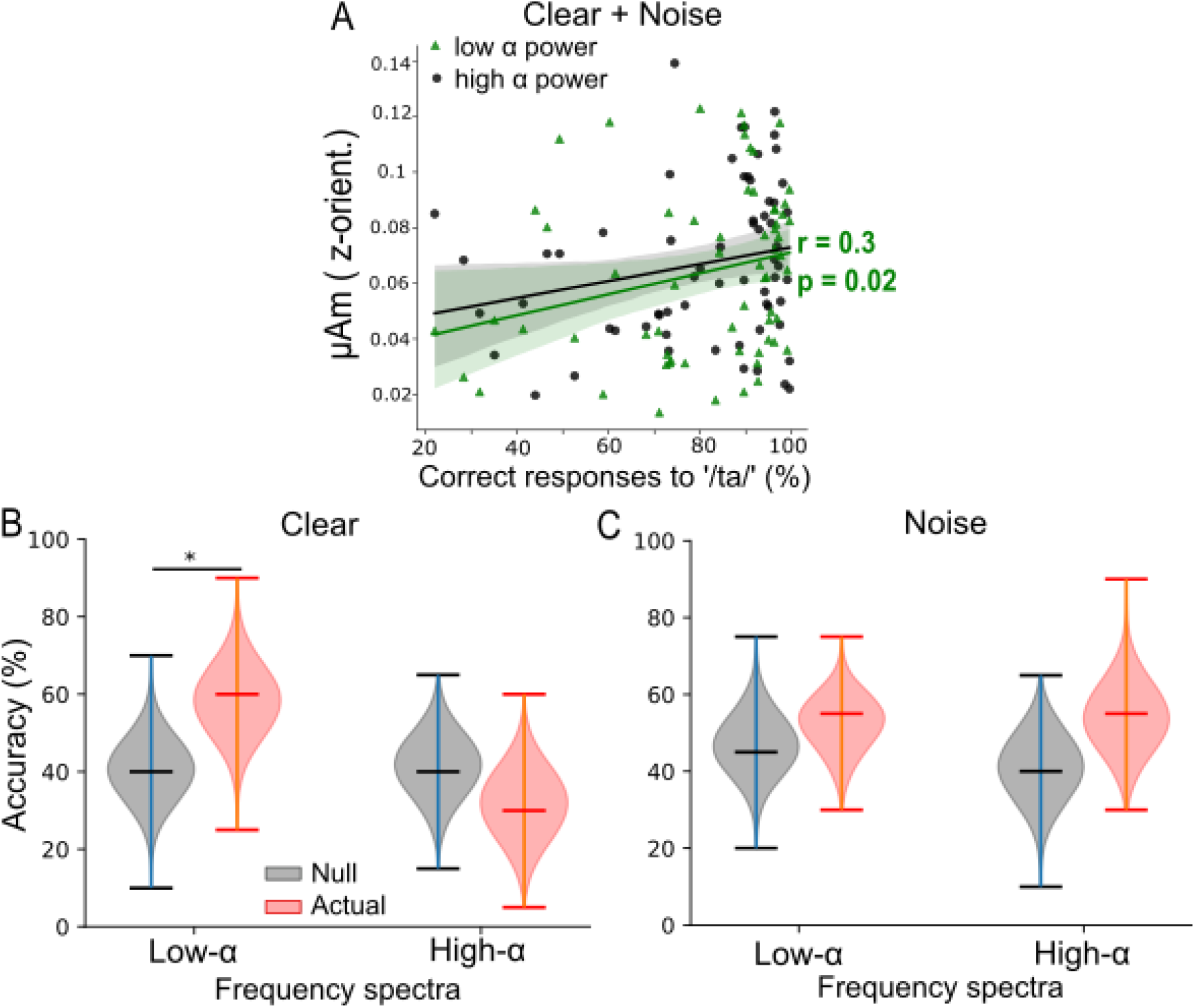
Low α FFRs better predict behavioral performance than high α FFRs during clear speech perception when pooling across background conditions. (A) Low α FFRs correlated positively with percent correct of speech target detection. r = Spearman's correlation. Shaded area=95% CI of the regression line. (B) For clear speech, SVM classifier accuracy was significantly better using low α (but not high α) FFR frequency spectra to classify subjects’ perceptual performance level (i.e., poor vs. good) compared to the null classification accuracies. (C) For noise-degraded speech, classification accuracies were similar when using low or high α FFR spectra and did not differ from the null classification accuracies. Upper/lower ticks=max/min; center tick=medians. * p< 0.01

### 3.4 Comparison of brain-behavior relations in the NH and HL groups

The aforementioned analyses showed that during high arousal states (i.e., with low α power), FFRs have a stronger relation with behavior compared to low arousal states, especially in the clear condition, when pooling the NH and HL groups. Hence, we studied the changes in low α FFR amplitudes with behavioral performance (QuickSiN, PTA, %-correct, and RTs) systematically by fitting linear regression models [Eq. (2)]. We observed significant coefficients or slopes (β) when PTA was used as *X* in Eq (2). For both the clear and noise-degraded speech, we found a significant negative slope between low α F0 amplitudes and PTAs in the NH group (β1 = −4.82×10^−9^, p = 0.01, Fig. 4A) but the slope of the HL group was not significantly different from the NH group. This indicates that even with clinically “normal” hearing, participants with slightly poorer thresholds have smaller FFRs during low α states. When separating the data by SNR, we observed similar trends of low α FFR amplitudes decreased with increased PTA in the NH and HL groups for clear speech (Fig. 4B) though the slopes were not significant. For noisy speech (Fig. 4C), we found that low α FFR amplitudes decreased significantly with increased PTA in the NH group (β1 = −6.53×10^−9^, p = 0.01) and the slope of the HL group was also significantly different (β3 = 6.77×10^−9^, p = 0.02) from the NH group. The fitted regression lines in Fig. 4C showed that low α FFR amplitudes were generally diminished by noise in the HL group.

**Figure 4.**
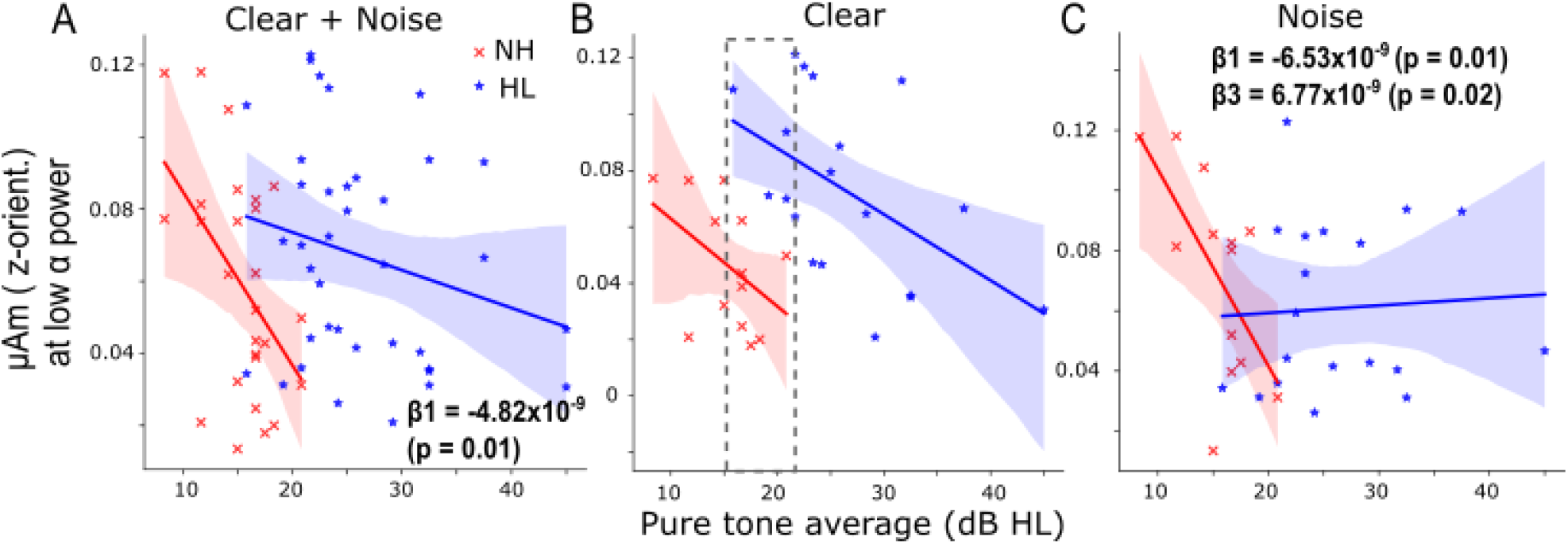
Hearing loss is associated with smaller speech-FFR amplitude under low cortical α (i.e., high arousal) states. (A) When pooling SNRs, the decrease (slope) of low α FFR F0 amplitudes as a function of PTA in the NH group was significant (β1 = −4.82×10^−9^, p = 0.01) and the slope of the HL group was not significantly different from the NH group. (B) For clear speech, similar trends of low α FFR amplitudes decreased with increased PTA were observed in the NH and HL groups though the slopes were not significant. The gray dashed box marks the overlapping PTA region for both groups. (C) For noise-degraded speech, the slope of low α FFR amplitudes decreased with increased PTA was significant (β1 = −6.53×10^−9^, p = 0.01) in the NH group and the slope of the HL group was significantly different (β3 = 6.77×10^−9^, p = 0.02) from the NH group. The fitted regression lines showed that low α FFR amplitudes were generally diminished by noise in the HL group. Shaded area=95 % CI of the regression line.

## 4 Discussion

Previous neuroimaging work reveals weaker functional connectivity between brainstem and cortex in older listeners with mild hearing loss than older adults with normal hearing for their age, and this interplay robustly predicts their SiN perceptual performance (Bidelman et al., 2019). Adding to these findings, we show the existence of active and dynamic modulation of brainstem speech processing in NH older listeners, which was dependent on online changes in listeners’ cortical state. This active and dynamic cortical-brainstem modulation, however, is diminished when processing speech in noise and in older adults with HL. Compared to NH listeners, HL listeners showed weaker parieto-occipital α power but those with minimal hearing loss (i.e., smaller PTA) had unusually large FFRs during low α states (gray dashed box in Fig. 4B). Although FFRs were smaller during low α power, they were predictive of perceptual speech measures (Fig. 3A) when pooling NH and HL participants and especially for clear speech (Fig. 3B). Collectively, our findings suggest that (i) FFRs during low α power (i.e., high cortical arousal states) are more predictive of behavioral performance, (ii) decreased low α FFRs with increased PTA in both NH and HL participants, and (iii) increased low α FFRs in older adults with mild hearing loss for clear speech suggesting an increase in central gain.

### 4.1 Effects of age on cortical α power and cortical modulation of brainstem speech processing

Cortical α indexes states of wakefulness and arousal (Pfurtscheller et al., 1996; Aftanas et al., 2002; Uusberg et al., 2013). Still, there is also evidence showing that α power may vary or index mind wandering during cognitive tasks (Compton et al., 2019; Maillet et al., 2020). Calculating the span length of low or high α trials for each listener showed averages of ~ 1.7 trials in both hearing groups, equating to several hundred milliseconds during our task. The relative speed of these fluctuations suggests that the α-modulations observed here are unlikely related to mind wandering *per se*, which presumably develops over longer time courses [tens of seconds (Pelagatti et al., 2020)]. Instead, we infer low α power tracks high arousal state while high α power reflects task focus but in a state of wakeful relaxation. Induced α activity is crucial for SiN perception as it might suppresses irrelevant information like noise to aid target speech processing (Strauß et al., 2014). In our previous study conducted in younger listeners (18-35 years) using similar EEG tasks, we observed larger α power to noise-degraded compared to clear speech during active engagement (see Fig. 2F in Lai et al., 2022). However, here in both NH and HL older adults, we do not find this same noise-related α effect. In general, α power was larger in NH than HL older listeners (Fig. 2C). We also observed that high α RMS values in young NH listeners (Fig. 2F in Lai et al., 2022) had a larger range than NH older adults. This observation advocates a reduction of α power with aging which has also been shown in other studies (Babiloni et al., 2006; Purdon et al., 2015). Decreased α activity is also related to declines in cognitive functions with increasing age (Klimesch, 1997, 1999).

More critically, we demonstrate the presence of dynamic and online modulation of brainstem speech encoding by fluctuations in cortical α state in older NH adults that are fundamentally different from those observed in younger, normal-hearing listeners (cf. Lai et al., 2022). In younger listeners, lower cortical α states positively correlate with smaller FFRs during SiN perception (Lai et al., 2022). Furthermore, low-α-indexed FFRs recorded in noisy backgrounds are predictive of behavioral RTs for rapid speech detection and have higher accuracies in token classification (Lai et al., 2022). Here, unlike younger listeners which require more difficult perceptual tasks (i.e., SiN perception) to tax the system and reveal effects of cortical arousal state on brainstem FFRs, we observed cortical modulation of FFRs in NH older adults during the perception of *clear* speech (Fig. 2D & E). Moreover, low-α-indexed FFRs associated more with behavior [speech detection (Fig. 3) and PTAs (Figs 4)] than high-α-indexed FFRs. Contrastively, in noise, low α and high α FFR amplitudes were not classifiable in terms of perceptual performance level. Taken together, the pattern of cortical-brainstem interactions in speech processing we found here in older NH listeners appears similar to what is found in younger listeners under challenging listening environments (cf. Lai et al, 2022). This indicates that aging might alter cross-talk between functional levels of the auditory system under challenging listening conditions as a means of compensatory processing. Similar maladaptive plasticity has been previously observed at higher cortical levels where frontal brain regions are more strongly engaged to aid auditory-sensory coding in superior temporal gyrus (Price et al., 2019). This further suggests the presence of age-related deficits in top-down modulation of brainstem speech processing by cortex and provides an explanation to why older listeners find it more exhausting to participate in cocktail party-like listening situations compared to younger listeners (Pichora-Fuller et al., 2017).

### 4.2 Effects of hearing loss on cortical α power and cortical modulation of brainstem speech processing

Compared to the NH group, we observed decreases in parieto-occipital α power in the HL group in both SNR conditions (Fig. 2C). Lower α power is reported in listeners with moderate hearing-loss across the age spectrum (Petersen et al., 2015). Moreover, PTA correlates negatively with pre-stimulus α power in older listeners (Alhanbali et al., 2021). These findings are partly concordant with our data since we found lower α power (during stimuli) in older listeners with mild to moderate hearing loss. Furthermore, in the HL group, we found no cortical-related enhancements of FFRs (i.e., F0 ratio ≈1) for clear speech, and responses were not different from the NH group in the noise condition though F0 ratio was > 1 (Fig. 2D). The interaction effect of SNR × α power was also distinct in direction between hearing groups. This finding implies that modulation of brainstem speech processing by cortical α state is altered in older listeners with mild hearing loss for both clear and noise-degraded speech processing.

In addition to parieto-occipital α power, trends of low-α-indexed FFRs to clear speech decreased with increased PTAs were observed in NH and HL listeners (Fig. 4B) and the slope of decrease was significant for noise-degraded speech in NH listeners (Fig 4C). The reduction in low α FFRs with poorer PTAs is probably related to the decrease in peripheral hearing ability. However, when comparing across groups at comparable hearing loss (PTA = 15-22 dB HL), we found enhanced FFR amplitudes in HL listeners (gray dashed box in Fig. 4B). Speculatively, this could indicate an increase in central gain, probably related to high-frequency (> 4 kHz) hearing loss in the HL group (our NH listeners had normal audiometric thresholds up to 4 kHz). Similar central gain compensation secondary to peripheral hearing loss has been observed previously in both animal and human neuroimaging studies (Bidelman et al., 2014; Chambers et al., 2016). These phenomena were completely collapsed by noise in the HL group where low α FFRs were relatively smaller in most HL listeners (Fig. 4C).

### 4.3 Association of brainstem speech processing during high arousal states with behaviors

In younger listeners and under noisy backgrounds, we previously showed that neural decoding applied to low α FFRs offered higher accuracies in token classification as compared to high α FFRs (see Fig. 5 in Lai et al., 2022). In this study, under no background noise, we observed frequency spectra of low α FFR had better classification accuracies for perceptual performance level than high α FFR. Thus, better speech token discrimination is consistently observed in FFRs during high arousal states in both younger and older adults. Furthermore, low α FFRs were also observed to be decreased as PTA increased, especially in the NH group for noise-degraded speech. These observations suggested that brainstem FFRs during high arousal states have a strong association with behavior perception.

## 5 Conclusion

Collectively, our study reveals age-related hearing loss not only reduces cortical α power but differentially alters its dynamic relationship with the subcortical speech processing especially in the presence of noise. While brainstem speech processing is actively modulated by cortical arousal state in normal-hearing older adults, this modulation is disrupted by signal degradations (i.e., noise) and hearing loss. Speech-FFRs during low α states also offer a higher fidelity representation of the acoustic speech signature and are more predictive of perceptual performance than FFRs yoked to states of high cortical α. Enhanced FFRs in older adults with near-normal hearing (i.e., very mild hearing loss) suggest the presence of increased central gain compensation for reduced auditory input (Bidelman et al., 2014; Chambers et al., 2016).

## 7 Acknowledgments

We would like to thank Dawei Shen and Stephen R. Arnott for their assistance with data collection as well as Yimei Li and Tushar Patni for providing consultation in statistical linear regression analysis.

