## Supplementary material for "Cortical-brainstem interplay during speech perception in older adults with and without hearing loss": Fig. 1 in Supplementary

### 1 Supplementary Figures

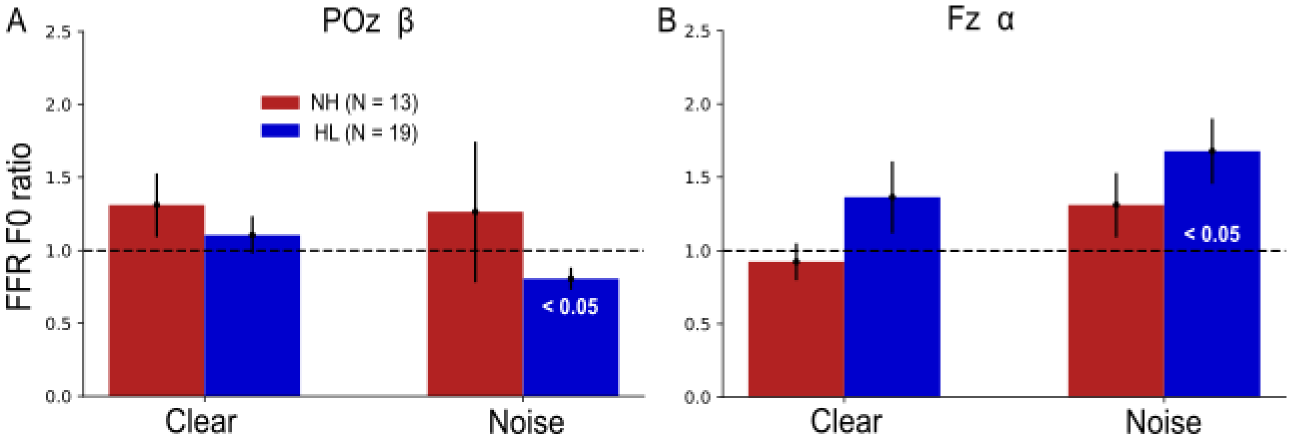


**Supplementary Figure 1.** The same FFR F0 ratio analysis was repeated for (A) POz α and (B) Fz α. FFR F0 ratios during low and high β or α trials were not significantly different between the NH vs. HL group in the clear and noise conditions. Only the FFR F0 ratios of the HL group in the noise condition (bar marked <0.05) are significantly different than 1 (1-sample t-test).
